# Re-evaluation of the report of *Culex quinquefasciatus* in the Republic of Korea: Molecular evidence of cryptic hybridisation within the *Culex pipiens* complex

**DOI:** 10.64898/2026.08.01.742187

**Authors:** Jiseung Jeon, Sunwoo Hwang, Sohyun Lee, Jihun Ryu, Kwang Shik Choi

## Abstract

The *Culex pipiens* complex comprises globally distributed mosquito species that serve as vectors for various arboviruses and filariasis. Female mosquitoes within the *Cx. pipiens* complex are morphologically very similar, making species identification based on morphologically characteristics challenging. To date, *Cx. pallens* and *Cx. pipiens* f. *molestus* have been reported to inhabit the Republic of Korea (ROK). Conversely, *Cx. quinquefasciatus*, predominantly distributed in tropical regions, has not been documented in the country since a single record in the 1950s. Recently, however, *Cx. quinquefasciatus* was reported from the Jeju region of the ROK based on molecular identification. This study aimed to re-evaluate the occurrence of *Cx. quinquefasciatus* in the ROK by conducting molecular identification using the *ace-2* and CQ11 markers on *Cx. pipiens* complex specimens collected nationwide, including from Jeju, where *Cx. quinquefasciatus* had previously been reported. Our analysis identified only *Cx. quinquefasciatus*–*Cx. pallens* hybrid individuals among the *Cx. pipiens* complex specimens from the Jeju region, highlighting the need to re-evaluate the analytical methods used in previous studies that reported the presence of pure *Cx. quinquefasciatus*. These findings provide essential evidence for future surveillance of the distribution, introduction, and establishment of *Cx. quinquefasciatus* and underscore the need for continuous monitoring.

## INTRODUCTION

A species complex refers to a group of closely related species that are difficult to distinguish morphologically, yet are genetically and ecologically distinct. Several mosquito species complexes have been identified, among which the *Culex pipiens* L. complex serves as a primary vector for disease-causing pathogens, such as West Nile virus and filarial parasites, and has a cosmopolitan distribution (Turell et al., 2001). The *Cx. pipiens* complex has long been regarded as a taxonomic challenge owing to minimal morphological variation across its members and complex life-history traits; however, recent advances in genomic technologies have begun to redefine its evolutionary history (Haba et al., 2025). Within the *Cx. pipiens* complex, *Cx. australicus* and *Cx. globocoxitus* inhabit Australia, whereas *Cx. quinquefasciatus* occurs predominantly in tropical regions, and *Cx. pipiens* is primarily found in temperate regions. Notably, both *Cx. quinquefasciatus* and *Cx. pipiens* exhibit global distribution patterns (Aardema et al., 2022). Furthermore, *Cx. pipiens* can be divided based on life-history traits into *Cx. pipiens* f. *pipiens* (bird-biting, diapausing in winter) and *Cx. pipiens* f. *molestus* (human-biting, autogenous) (Haba and McBride, 2022). *Culex pallens* (previously known as *Cx. pipiens pallens*) was recently elevated to full species status and is known to occur primarily in East Asia (Aardema, 2022; Harbach, 2023). The distribution and density of these species are expected to change as a result of climate change, which is anticipated to have a particularly significant impact on the geographic range of *Cx. quinquefasciatus*, a species of tropical origin (Samy et al., 2016). Consequently, early detection of the invasion of major disease vectors, alongside continuous surveillance, is paramount for suppressing outbreaks of vector-borne diseases (Giunti et al, 2023).

Traditionally, the Republic of Korea (ROK) has been classified as a temperate region; however, recent climate change has accelerated subtropicalisation across the southern regions of the Korean Peninsula, including Jeju Island (Rahman et al., 2025). Although the presence of *Cx. quinquefasciatus* was recorded in the ROK in the 1950s (Chu, 1956), no official records of its occurrence have been reported since. Nationwide surveys employing molecular markers have revealed that only two species of the *Cx. Pipiens* complex, *Cx. pipiens* f. *molestus* and *Cx. pallens*, currently inhabit the ROK (Ryu and Choi, 2022). Furthermore, Ryu and Choi (2022) conducted molecular identification of specimens collected in Jeju in 2020; however, *Cx. quinquefasciatus* was not detected. Recently, Kwon et al. (2025) reported the occurrence of *Cx. quinquefasciatus* for the first time based on molecular identification of *Cx. pipiens* complex specimens.

Currently, identification of the *Cx. pipiens* complex is commonly conducted using a polymerase chain reaction (PCR) assay that exploits species-specific polymorphisms in the acetylcholinesterase-2 (*ace-2*) gene (Smith and Fonseca, 2004; Ryu and Choi, 2022). However, the *ace-2*-based assay cannot differentiate between the two ecological forms of *Cx. pipiens*. Consequently, an additional molecular assay targeting the CQ11 locus is required to distinguish *Cx. pipiens* f. *molestus* from *Cx. pipiens* f. *pipiens* (Bahnck and Fonseca, 2006). To date, no study in the ROK has integrated both the ace-2 gene and the CQ11 locus for identifying the *Cx. pipiens* complex. Because members within the *Cx. pipiens* complex exhibit distinct life-history traits and vector competence (Farajollahi et al., 2011; Turell et al., 2011; Haba and McBride, 2022), accurate species identification and tailored control strategies are essential for efficient vector management.

Hence, the present study was undertaken to address the following two critical questions: (i) given that *Cx. quinquefasciatus* was not detected in the 2020 Jeju survey but was newly reported in the 2025 survey, was this species truly absent from Jeju prior to 2025? (ii) Given the traditional use of the *ace-2*-based PCR assay in the ROK—which is insufficient to differentiate between the two ecotypes of *Cx. pipiens*—does *Cx. pipiens* f. *molestus* indeed occur exclusively within the country? To address these questions, this study conducted molecular identification using both *ace-2* and CQ11 markers for *Cx. pipiens* complex specimens collected nationwide in the ROK from 2020 to 2026. The findings of this study elucidate the distribution patterns of the *Cx. pipiens* complex, a primary group of disease-vector mosquitoes, and provide fundamental insights that can inform vector surveillance and management strategies in response to future climate change.

## MATERIALS AND METHODS

### Sample collection

To collect specimens of the *Cx. pipiens* complex, sampling was conducted from 2020 to 2026 across 17 locations, targeting residential areas and cattle sheds throughout the ROK. Female mosquitoes were captured using BG-Sentinel traps (Biogents, Regensburg, Germany) and black light traps (BT Global, Seongnam, ROK), supplemented with dry ice as an attractant. To evaluate the applicability of the molecular markers to male specimens, male individuals were also included in this study. Male *Cx. pallens* were collected from mating swarms using the sweeping method, whereas male *Cx. pipiens* f. *molestus* were obtained from colonies maintained in the insectary (Animal Systematics and Taxonomy Laboratory, Kyungpook National University, ROK).

The collected mosquitoes were transported to Kyungpook National University (Daegu, ROK) and stored at −20 °C prior to species identification to prevent sample degradation. Specimens belonging to the *Cx. pipiens* complex were distinguished from other mosquito species using morphological keys. Subsequently, genomic DNA (gDNA) was extracted from *Cx. pipiens* complex individuals using the Clear-S™ Quick DNA Extraction Kit (InVirusTech, Gwangju, ROK).

### Molecular identification

To identify members of the *Cx. pipiens* complex, multiplex PCR assays were performed using gDNA. First, molecular identification based on the *ace-2* region was conducted to differentiate among *Cx. pallens*, *Cx. pipiens*, and *Cx. quinquefasciatus* within the *Cx. pipiens* complex (Ryu and Choi, 2022). The primers used in this study (F1457: 5′-GAG GAG ATG TGG AAT CCC AA-3′; ACEmole_R: 5′-TTC TCA CAG AGC CAT CAT CGA C-3′; ACEpall_R: 5′-ACA TGT CAA AAG CTC AGT TAG T-3′; and ACEquin_R: 5′-TGC CAC AGC CAT TCA AGA AGG-3′) were designed to generate species-specific amplification products (*Cx. pipiens*: 183 bp; *Cx. pallens:* 291 bp; *Cx. quinquefasciatus*: 449 bp) (**Fig. 1a**). The PCR conditions were as follows: each PCR reaction mixture had a total volume of 25 μL and contained 1 μL of extracted gDNA template, 1× PCR buffer, 0.2 mM dNTPs, 0.4 μM of each primer, and 0.5 units of Taq DNA polymerase. The PCR cycling conditions consisted of an initial denaturation at 94 °C for 5 min, followed by 35 cycles of 94 °C for 30 s, 56 °C for 30 s, and 72 °C for 30 s, followed by a final extension at 72 °C for 5 min.

**Fig. 1.**
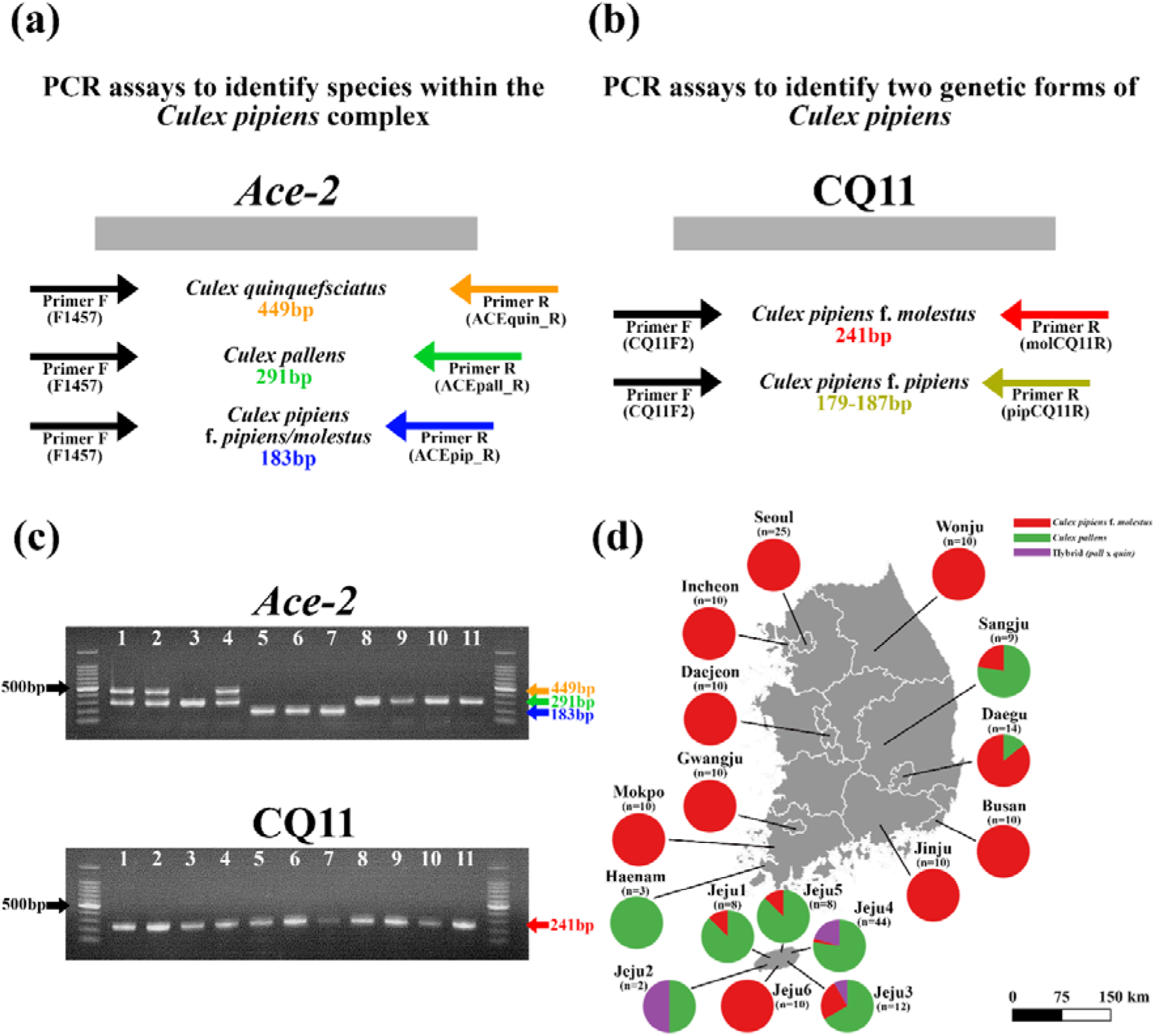
Molecular identification of the *Cx. pipiens* complex based on the *ace-2* gene (**a**). Form-specific identification based on the CQ11 locus (**b**). Representative PCR gel electrophoresis results using *ace-2* and CQ11 markers: *ace-2* assay [lanes 1–2: *Cx. quinquefasciatus–Cx. pallens* hybrid (female); lanes 3, 5–7: *Cx. pallens* (female); lane 4: *Cx. pallens* (male); lanes 8–10: *Cx. pipiens* (female); lane 11: *Cx. pipiens* (male)] and CQ11 assay [lanes 1–11: *Cx. pipiens* f. *molestus*] (**c**). Geographic distribution of *Cx. pipiens* complex species collected across sampling sites in this study (**d**).

Subsequently, for specimens identified as *Cx. pipiens* based on the *ace-2* region, molecular identification was performed using the CQ11 locus to distinguish between *Cx. pipiens* f. *molestus* and *Cx. pipiens* f. *pipiens* (Bahnck and Fonseca, 2006). The primers used (CQ11F2: 5′-GAT CCT AGC AAG CGA GAA C-3′; pipCQ11R: 5′-CAT GTT GAG CTT CGG TGA A-3′; and molCQ11R: 5′-CCC TCC AGT AAG GTA TCA AC C-3′) were designed to yield distinct amplification products for each ecological form (*Cx. pipiens* f. *molestus*: 241 bp; *Cx. pipiens* f. *pipiens*: 179–187 bp) (**Fig. 1b**). The PCR conditions were as follows: each PCR reaction mixture had a total volume of 25 μL and contained 1 μL of extracted gDNA template, 1× PCR buffer, 0.2 mM dNTPs, 0.4 μM of each primer, and 0.5 units of Taq DNA polymerase. The PCR cycling conditions comprised an initial denaturation at 94 °C for 5 min, followed by 35 cycles of 94 °C for 30 s, 54 °C for 30 s, and 72 °C for 30 s, with a final extension at 72 °C for 5 min.

All PCR amplification products were resolved on 1.5% agarose gel, and bidirectional sequencing was performed using the respective PCR primers (Macrogen, Daejeon, ROK). All sequence data obtained in this study have been deposited in NCBI GenBank (accession numbers: PZ764115–PZ764131).

### Data analysis

Phylogenetic analysis of the *ace-2* sequences obtained from *Cx. pipiens* complex individuals was conducted. Sequence alignment was performed using the L-INS-i method in MAFFT v.7 (Katoh et al., 2019). W-IQ-TREE was used for maximum likelihood (ML) analysis and substitution model selection (Trifinopoulos et al., 2016). Based on the Bayesian Information Criterion (BIC), the substitution model for each region was determined to be F81+F, and 1,000 bootstrap replicates were applied to evaluate node support. The phylogenetic tree was visualised using FigTree v.1.4.4 (http://tree.bio.ed.ac.uk/software/figtree/).

For the site where *Cx. quinquefasciatus* was confirmed, backward trajectory analysis was performed to estimate the introduction route and point of origin based on airflow dynamics. NOAA’s Hybrid Single-Particle Lagrangian Integrated Trajectory (HYSPLIT) model was used for the analysis (Stein et al., 2015). The backward trajectory analysis was conducted using the web-based version (https://www.arl.noaa.gov/hysplit/) with Global Data Assimilation System (GDAS) meteorological data. The arrival height was set to 500 m above ground level. The analysis covered a period of 300 h prior to the collection date *of Cx. quinquefasciatus*, with trajectories calculated at 12-h intervals.

## RESULTS AND DISCUSSION

### Molecular identification of the *Culex pipiens* complex

In this study, molecular identification was conducted on 205 specimens of the *Cx. pipiens* complex collected across the ROK. For specimens identified as *Cx. pipiens* based on the *ace-2* region, further identification of the two ecological forms was performed using the CQ11 locus (**Fig. 1c**). Among the 205 specimens analysed, 125 were identified as *Cx. pipiens* f. *molestus*, 69 as *Cx. pallens*, and 11 as *Cx. quinquefasciatus*–*Cx. pallens* hybrid individuals (**Fig. 1d**). Analysis using the CQ11 locus revealed no *Cx. pipiens* f. *pipiens* individuals, indicating that all specimens belonged to *Cx. pipiens* f. *molestus*. The *Cx. quinquefasciatus–Cx. pallens* hybrid individuals were detected exclusively on Jeju Island, with 9 of the 11 specimens collected from the Jeju4 site (Jeju2: 1, Jeju3: 1, Jeju4: 9). No pure *Cx. quinquefasciatus* specimens were detected. The hybrid individuals were confirmed in 2023 (n = 8) and 2024 (n = 3), indicating that this lineage may have been introduced to Jeju Island prior to the initial report in 2025.

Consistent with previous findings, a double band corresponding *to Cx. pallens* (291 bp) and *Cx. quinquefasciatus* (449 bp) was observed in *Cx. pallens* males (Fonseca et al., 2009; Ryu and Choi, 2022). This phenomenon is likely attributable to the physical linkage of the *ace* locus to the male-determining locus (MDL), leading to sex-linked asymmetric introgression from *Cx. quinquefasciatus* into *Cx. pallens* males (Fonseca et al., 2009). These results are further supported by the phylogenetic analysis of the *ace-2* gene using sequences obtained from the *Cx. quinquefasciatus–Cx. pallens* hybrid specimens and *Cx. pallens*, in which sequences derived from the hybrids and *Cx. pallens* males clustered within a single clade (**Fig. 2a**). Although *Cx. pallens* is known to be restricted to East Asia, including the ROK, Japan, and China, its evolutionary origin remains unclear (Aardema et al., 2022). Given the limited dataset obtained in the present study, it is difficult to determine precisely when this introgression event occurred in *Cx. pallens* males and whether it is ongoing. Therefore, further genomic-level studies are warranted to elucidate the evolutionary origin of *Cx. pallens*.

**Fig. 2.**
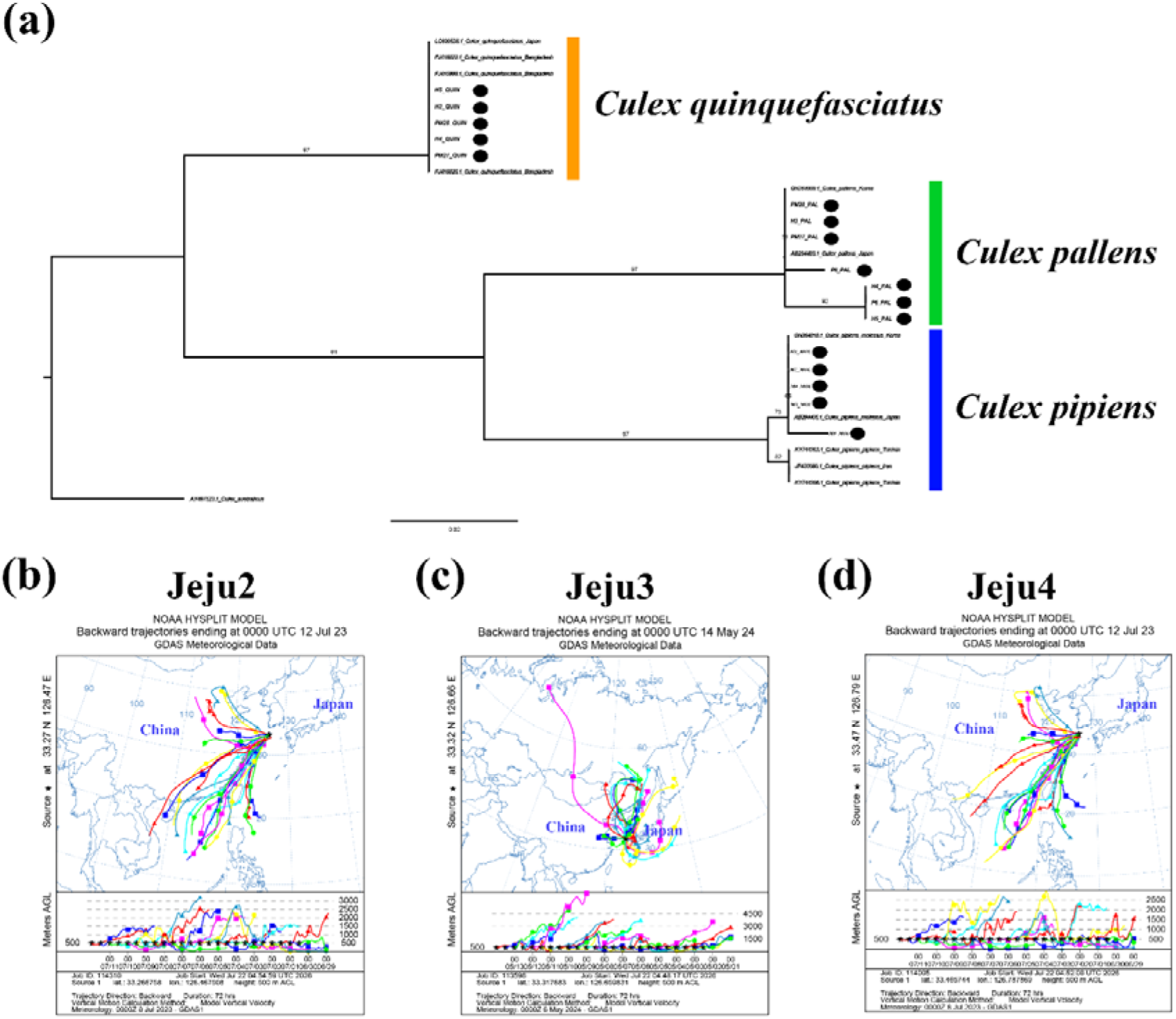
Phylogenetic tree based on *ace-2* gene sequences. Bootstrap values based on 1,000 replicates are shown at the nodes. Sequences retrieved from NCBI GenBank are denoted by their accession numbers, and black circles indicate samples obtained in the present study (H: hybrid; PM: *Cx. pallens* male; M: *Cx. pipiens* f. *molestus*) **(a)**. Back trajectory analysis showing air-mass movement toward the sampling sites where *Cx. quinquefasciatus*–*Cx. pallens* hybrids were collected: Jeju2 **(b)**, Jeju3 **(c)**, and Jeju4 **(d)**. Each line represents a distinct trajectory. The top portion of each image displays a horizontal path, whereas the bottom portion illustrates the elevation of the path.

Kwon et al. (2025) performed molecular identification on *Cx. pipiens* complex specimens collected from Jeju in 2025, reporting the presence of both pure *Cx. quinquefasciatus* and *Cx. pipiens–Cx. quinquefasciatus* hybrid individuals. However, their identification methodology warrants careful reconsideration. Although they employed the *ace-2* locus-based PCR assay developed by Smith and Fonseca (2004), they incorporated only two primers (*Cx. pipiens* and *Cx. quinquefasciatus*) out of the four primers originally designed to identify the *Cx. pipiens* complex (*Cx. pipiens, Cx. quinquefasciatus, Cx. pallens*, and *Cx. australicus*) (Kwon et al., 2025). Consequently, they reported that pure *Cx. quinquefasciatus* was present in the ROK. Nevertheless, because the primer specific for *Cx. pallens* was omitted from their molecular assay, any *Cx. quinquefasciatus*–*Cx. pallens* hybrid specimens analysed using this assay would potentially be misidentified as pure *Cx. quinquefasciatus*. Given that no pure *Cx. quinquefasciatus* were detected in the present study, alongside the omission of the *Cx. pallens*-specific primer in the previous molecular assay, the assertion that pure *Cx. quinquefasciatus* is currently present in the ROK requires further verification.

To investigate the possibility of long-distance windborne influx from overseas via airflow dynamics, backward trajectory analysis was conducted for approximately two weeks prior to the collection dates at the three sites where hybrid individuals were collected (Jeju 2–4). The results revealed that Jeju 2 and 4 displayed similar airflow trajectory patterns, whereas no significant airflow pattern was observed for Jeju 3 (**Fig. 2b–d**). For Jeju 2 and 4, the majority of air masses originated from the eastern and southern coastal regions of China, areas where *Cx. quinquefasciatus* is widely distributed (Liu, 2020). Considering that overwintering in the ROK is challenging for *Cx. quinquefasciatus* given its tropical origin and the ROK’s low average winter temperatures, along with the atmospheric characteristics of Jeju Island situated within the prevailing westerlies belt, the hybrid individuals identified in the ROK are likely attributable to windborne influx from China. Such windborne migration from China to the ROK has been well documented for major agricultural pests such as the brown planthopper (*Nilaparvata lugens*) (Jeong et al., 2024), and similar cases have been reported for *Cx. tritaeniorhynchus*, a primary vector of the Japanese encephalitis virus (Jeon et al., 2024). The novel emergence of closely related species within the *Cx. pipiens* complex that were not previously established poses a risk beyond the introduction of a disease vector; it may serve as a conduit for the dissemination of genes associated with vector control, such as insecticide resistance, through hybridisation. Currently, rapid subtropicalisation is underway across the southern regions of the ROK owing to climate change. Therefore, subsequent studies on population genetics are urgently required to validate both the overseas windborne influx hypothesis proposed in this study and the potential establishment of *Cx. quinquefasciatus*.

### Cryptic hybridisation within the *Culex pipiens* complex

As previously mentioned, current evidence regarding the presence of pure *Cx. quinquefasciatus* in the ROK remains insufficient; rather, it is highly likely that *Cx. quinquefasciatus–Cx. pallens* hybrids identified in this study, along with the *Cx. quinquefasciatus–Cx. pipiens* hybrids reported by Kwon et al. (2025), are currently distributed in the country. Three non-mutually exclusive hypotheses may account for these observations: (i) a small number of pure *Cx. quinquefasciatus* entered the ROK and subsequently hybridized with indigenous *Cx. pipiens* complex populations; (ii) hybrid individuals, rather than pure individuals, were introduced from the outset; or (iii) both mechanisms have contributed to the present scenario. Because the winter climate of temperate ROK poses a significant barrier to the overwintering of tropical-origin *Cx. quinquefasciatus*, pure individuals, even if introduced, may have failed to overwinter. Consequently, only hybrid populations that acquired cold-tolerance traits through hybridisation with *Cx. pipiens* or *Cx. pallens* might have survived the Korean winter and become established. Furthermore, given that *Cx. quinquefasciatus* and *Cx. pallens* occur sympatrically in eastern and southern China—the potential origins of influx—forming a hybrid zone (Liu et al., 2020), the possibility that only lineages capable of adapting to temperate climates have been introduced and established cannot be ruled out. Notably, a high hybridisation rate between *Cx. pallens* and *Cx. quinquefasciatus* populations was previously reported in Saga, Japan (Fonseca et al., 2009), a region situated at a latitude similar to Jeju, where hybrid specimens were obtained in this study. Nevertheless, the fitness costs or benefits that such hybridisation imparts to the adaptability of *Cx. pipiens* complex populations remain unclear, highlighting the need for further investigation.

To date, studies investigating the *Cx. pipiens* complex using molecular markers in the ROK remain remarkably limited (Ryu and Choi, 2022; Kwon et al., 2025). Particularly, because the *Cx. pipiens* complex exhibits high interspecific hybridisation rates accompanied by frequent backcrossing, single molecular markers are insufficient for definitive species identification (Cornel et al., 2012; Ohashi et al., 2014). Given the limited sample size analysed in the present study, continuous surveillance is warranted to monitor the prevalence of hybrids and the potential establishment of *Cx. quinquefasciatus* in both Jeju and mainland regions. Such efforts will provide an exemplary case study regarding the suppression of vector-borne disease outbreaks following novel introductions, as well as the influx and establishment of disease vectors driven by climate change.

## Acknowledgements

The authors thank the Korea Disease Control and Prevention Agency (KDCA) for financial support.

## Funding

This work was supported by the Korea Disease Control and Prevention Agency (grant code: 6332-305-320).

### Data availability

All data generated or analysed during this study are included in this published article.

## Author contributions

**Jiseung Jeon**: Conceptualization; investigation; methodology; formal analysis; validation; data curation; writing – original draft; review and editing. **Sunwoo Hwang**: investigation; formal analysis; validation. **Sohyun Lee**: investigation; formal analysis; validation. **Jihun Ryu**: investigation; formal analysis; validation**. Kwang Shik Choi:** Conceptualization; funding acquisition, supervision; project administration; resources; review and editing.

## Declarations

### Ethics approval and consent to participate

Not applicable.

### Consent for publication

Not applicable.

### Competing interests

The authors declare that they have no competing interests.

## Notes

### Competing Interest Statement

The authors have declared no competing interest.

